# Non-High Frequency Heart Rate Chaos: A Noninvasive Marker of REM Sleep and Obstructive Sleep Apnea Syndrome in Children

**DOI:** 10.1101/457630

**Authors:** Zhi-De Deng, Natalia M. Arzeno, Eliot S. Katz, Helena Chang, Carole L. Marcus, Chi-Sang Poon

## Abstract

Obstructive sleep apnea syndrome (OSAS) is a highly prevalent condition associated with considerable metabolic, cardiovascular, and neurocognitive morbidity. Childhood OSAS is underdiagnosed due to a limited number of sleep laboratories and the lack of a screening test, and the subtlety of daytime symptoms in children compared to adults. A potential marker of OSAS is apnea-induced sympathoexcitation, which is likely to be exacerbated during rapid-eyemovement (REM) sleep. However, traditional methods of assessing sympathetic activity are either too invasive or insensitive/nonspecific for clinical use, particularly as a screening test. Study population comprised pediatric patients with OSAS (16 moderate/severe, 18 mild) and 18 normal non-snoring controls. We show that the chaotic dynamics of heart rate variability (HRV) as assessed by a sensitive noise titration assay is significantly increased during REM compared to non-REM sleep in children, particularly those with OSAS. The increase in heart rate chaos prevails in the face of decreased parasympathetic-mediated high-frequency component of the HRV power spectrum, indicating that the chaos was correlated to sympathetic instead of parasympathetic activity. Receiver operating characteristic analysis shows that such non-high frequency chaos reveals changing sympathetic–parasympathetic activities that are not discernible by conventional HRV metrics such as low- to high-frequency power ratio or sample entropy, with sensitivity and specificity sufficient to detect even mild OSAS in children. Results suggest a possible role for non-high frequency heart rate chaos as a selective noninvasive marker of sympathoexcitation in REM sleep, OSAS and potentially other cardiovascular abnormalities such as congestive heart failure.

## I. Introduction

Obstructive sleep apnea syndrome (OSAS) is characterized by intermittent episodes of disrupted breathing due to pharyngeal narrowing or collapse, resulting in hypoxemia, hypercapnia, and/or sleep disruption. OSAS reportedly affects at least 2% of children in the United States and Europe, though about 10% habitually snore [1]. The etiology and clinical manifestations of OSAS in children are quite different from adults [1–5]. In particular, children with OSAS may have normal sleep stage distribution, few electrocortical arousals, and obstructive events occurring predominantly during rapid eye movement (REM) sleep [6].

Airflow obstruction and REM sleep may both exert profound influences on autonomic regulation. Sympathetic nerve activity typically increases during REM sleep [7] and with OSAS in adults [8–10]. These effects are accompanied by concurrent parasympathetic withdrawal as identified from power spectrum analysis of heart rate variability (HRV), a widely used noninvasive method of assessing beat-to-beat cardiac-autonomic regulation [11, 12]. It has been reported that the HRV power spectrum in children has a significantly decreased high-frequency (HF) component and increased low- to high-frequency power ratio (LF/HF) during REM compared to non-REM (NREM) sleep and with moderate/severe OSAS compared to normal controls, indicating downregulation of parasympathetic activity in these conditions [13–15]. However, since the LF power and LF/HF are nonspecific indicators of sympathetic outflow (which is a major drawback of conventional HRV analyses [12, 16–18]), it remains uncertain whether sympathetic activity is upregulated in children during REM sleep or with mild OSAS. A noninvasive method with high sensitivity and specificity in detecting even mild OSAS and associated autonomic abnormalities is critical in developing an effective home screening test for early diagnosis [19, 20], as even mild OSAS may be associated with considerable neurocognitive morbidity in children despite the subtlety of daytime symptoms [21].

An emerging approach to assess cardiac-autonomic modulation of HRV is based on methods of nonlinear time series analysis. Nonlinear control of HRV has been demonstrated in infants [22, 23]. Recent studies using a sensitive noise titration assay of nonlinear dynamics [24, 25] have shown that HRV in healthy young and elderly subjects has distinct chaotic signatures as measured by the noise limit (NL, see Methods and Supplementary Information), which can be correlated positively with the HF component and negatively with LF/HF particularly during nighttime (i.e., HF chaos) [26, 27]. Here, we show that a novel form of heart rate chaos called non-HF chaos (i.e., independent of the HF component) provides a more robust measure of changing cardiac sympathetic-parasympathetic activities than is possible with conventional HRV metrics, with sensitivity and specificity sufficient to detect even mild OSAS in children.

## II. Methods

### A. Subjects

The subjects enrolled in this study underwent a comprehensive overnight polysomnogram evaluation of obstructive apnea as part of the research protocols approved by the Institutional Review Boards and reported previously [28–30]. Obstructive apnea is defined as cessation of oronasal airflow in presence of respiratory efforts for at least two respiratory cycle times [2]. Obstructive hypopnea is defined as a reduction in amplitude of oronasal airflow (≥ 50%) accompanied by a 4% oxygen desaturation and/or arousal. The apnea–hypopnea index (AHI) is defined as the number of obstructive apnea and hypopnea events per hour of sleep. OSAS was classified into two severity levels: mild (AHI = 1 to 5/hour) and moderate/severe (AHI > 5/hour) [31].

The study population comprised 52 children (age 1–16 yr), of which 16 were classified as moderate/severe OSAS, 18 were mild/borderline OSAS, and 18 were normal; corresponding demographic and polysomnographic data are given in Table 1. The normal group included 10 children recruited from the community for the study and had no history of sleep disorders, and eight others who presented to the clinic with symptoms sufficient to warrant a sleep study, which subsequently showed that they neither snored, nor had apnea. Informed consent was obtained from the parent and assent from children more than 5 years of age. All subjects were free of lung or neuromuscular disease, cardiac pathology or arrhythmia.

**Table 1:** Demographic and polysomnographic data

|  | Non-snoring | Mild OSAS | Moderate/severe OSAS |
| --- | --- | --- | --- |
| *N* | 18 | 18 | 16 |
| Male (*N*) | 10 | 13 | 12 |
| Age (years, range) | 8.0 ± 4.8 (2–16) | 4.7 ± 3.5 (1–13) * | 4.8 ± 2.2 (2–8) * |
| HR (bpm) | 80.3 ± 12.0 | 82.5 ± 7.4 | 95.9 ± 17.6 |
| SWS (%TST) | 26.3 ± 11.5 | 24.8 ± 10.2 | 27.4 ± 6.3 |
| REM (%TST) | 19.5 ± 5.8 | 20.6 ± 5.8 | 20.5 ± 6.7 |
| AHI (Events/hr) | 0.2 ± 0.3 | 2.1 ± 1.3 * | 21.6 ± 20.5 * |
| SpO<sub>2</sub> Nadir (%) | 94.2 ± 3.8 | 91.8 ± 3.6 | 79.6 ± 15 * |
| Arousal index (Events/hr) | 7.2 ± 2.5 | 8.8 ± 2.3 | 13.7 ± 7.4 * |

Data are presented as mean ± SD. HR, mean heart rate; SWS, slow wave sleep; REM, rapid eye movement; %TST, percent of total sleep time; AHI, apnea–hypopnea index; SpO<sub>2</sub>, arterial oxygen saturation; OSAS, obstructive sleep apnea syndrome.
\* Indicates significant difference compared to non-snoring controls ( $p < 0.05$ ).

### B. Sleep and ECG Data

Sleep stages were scored in 30-second epochs. For each subject, we selected for analysis a continuous 3-hour ECG recording in the middle of the night that demonstrated minimal missing data and movement artifacts. In preliminary analyses we found no significant difference between NL values in stage 2 and slow-wave sleep. Accordingly, all data in these sleep stages were combined as NREM sleep. The 3-hour recording included at least one continuous 10-minute segment each of REM and NREM sleep. To minimize selection bias, the 3-hour recordings were carefully selected to capture apnea/hypopnea events consistent with the all-night AHI of each subject.

The ECG was digitized at a sampling rate of 100 Hz. The R-waves’ fiducial points were detected using a Hilbert transform-based peak extraction algorithm [32]. The R-wave to R-wave interval (RRI) time series was derived by calculating the sequential intervals between consecutive QRS peaks. The RRI was visually inspected for artifacts, and ECG segments that were clearly nonphysiological were removed. No further attempt was made to distinguish normal sinus beats from ectopic beats as the latter are rare among children in this age range [33] and the artificial elimination of such suspected ectopic beats could itself introduce artifacts [34], particularly during apnea events. The RRI time series was converted into equally-spaced samples in time by cubic-spline interpolation with a sampling rate of 4 Hz. All data manipulation and analysis algorithms were implemented in MATLAB (The MathWorks, Inc., Natick, MA). The RRI series were analyzed within a 5-minute time window that slid at 30-second intervals. An example patient sleep stages and RRI times series is illustrated in Figure 1.

**Figure 1:**
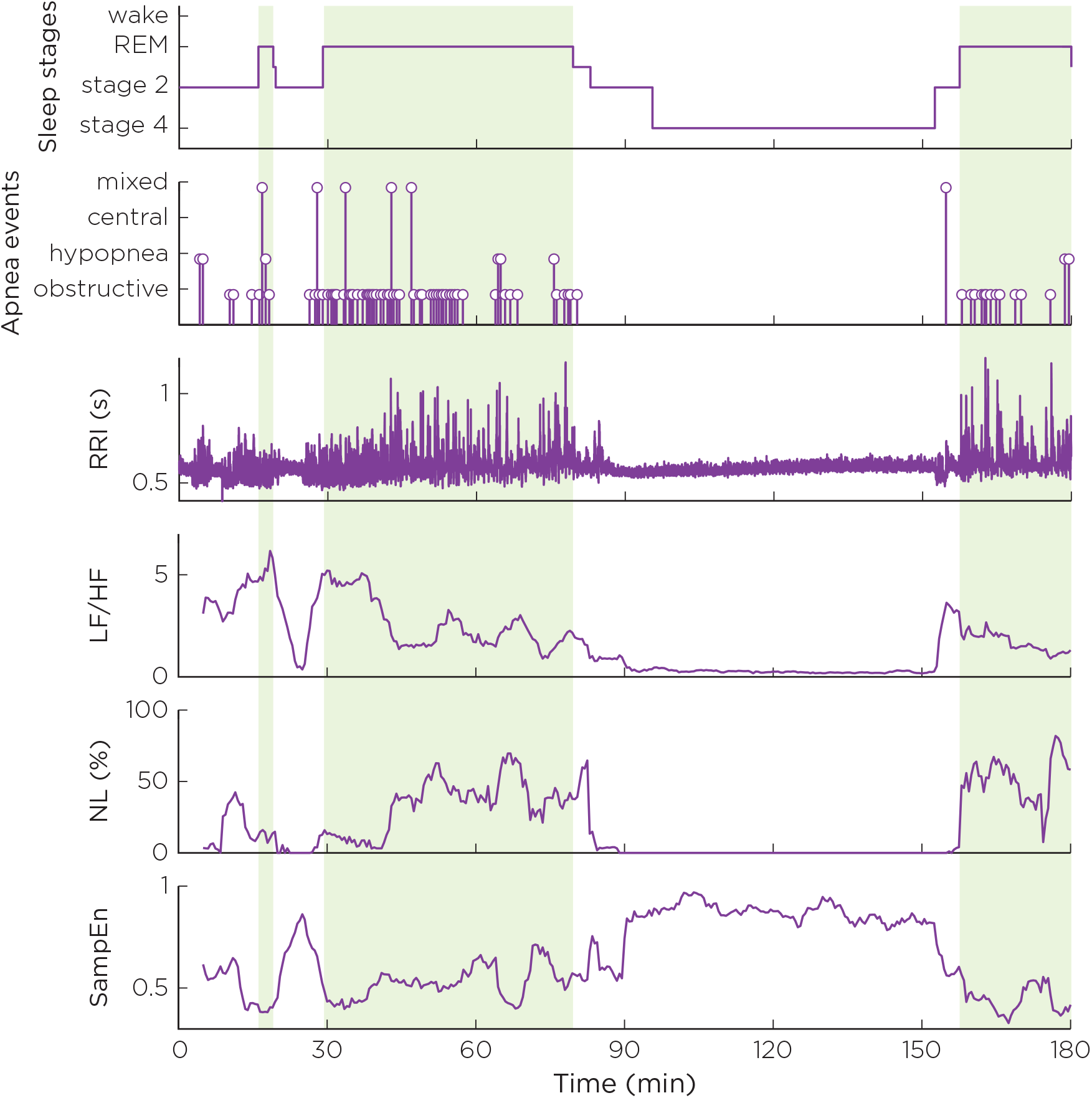
Time series of selected HRV metrics during sleep for a patient with moderate/severe OSAS. The rows are (from top to bottom): sleep stages, with REM sleep epochs shaded; time stamps of respiratory events; RRI time series; low-frequency to high-frequency (LF/HF) power ratio; noise limit (NL); and sample entropy (SampEn).

### C. HRV Analysis

#### 1) Time-Domain Analysis

Five common time-domain HRV indices were evaluated [11]: SDNN, SDANN, RMSSD, pNN50, and triangular index. Each index was calculated both over the entire RRI series and over all the REM and NREM sub-segments in the RRI series.

#### 2) Frequency-Domain Analysisx

Power spectrum of HRV was estimated by using the discrete pseudo Wigner–Ville distribution (PWVD), which allows continuous tracking of the changes in frequency and amplitude of each spectral component in nonstationary (apnea-prone) RRI series [35]. The PWVD partially suppresses cross-term interference of the Wigner–Ville transform through frequency smoothing. No additional time smoothing was performed in order to preserve temporal resolution. Results of the pseudo Wigner–Ville algorithm were also verified against the conventional Welch periodogram method. From each spectrum, we calculated the power in two relevant frequency bands according to established guidelines [11]: low frequency (LF), from 0.04 Hz to 0.15 Hz; and high frequency (HF), from 0.15 Hz to 0.4 Hz. Because normalized LF and HF indices are mathematically redundant [12, 36], only the absolute powers and the low- to high-frequency power ratio (LF/HF) are presented in this study. We found that the LF and HF powers demonstrated log-normal distributions; hence the log transformed values [ln(LF) and ln(HF)] were analyzed instead.

#### 3) Entropy Analysis

Sample entropy (SampEn), a refinement of approximate entropy [37], was computed to assess the statistical complexity/regularity of the HRV data. The SampEn algorithm computes the likelihood that epochs of length *m* that are similar within a tolerance *r* remain similar for epochs of length *m* +1. We chose *m* =2 and *r* = 20% of the standard deviation for the 5-minute segment.

#### 4) Noise Titration Analysis

The noise titration technique [24, 25] was used to assess changes in chaotic dynamics in noise-contaminated RRI time series (see Supplementary Information). The resulting titration index, the noise limit (NL), provides a measure of the relative level of chaos in HRV within each time window.

### D. Statistical Analysis

Possible interactions of OSAS and sleep effects on the HRV metrics were assessed based on a linear mixed-effects model with a random subject effect using OSAS severity (3 levels: moderate/severe OSAS, mild OSAS, and non-snoring control) and sleep states (2 levels: REM and NREM) as independent variables. The linear mixed-effects model was also used to account for correlations between repeated measures of HRV metrics within the same individual [38]. Post-hoc pairwise comparisons between patient groups were performed using a Bonferroni correction. Receiver operating characteristic (ROC) was used to assess the discriminatory power of selected HRV metrics in detecting OSAS based on heart rate data alone. The area under the ROC curve (AUC) was used to assess test performance. All statistical analyses were performed using SAS 9.2 (SAS Institute, Inc., Cary, NC).

## III. Results

### A. NL is Increased in REM Sleep and OSAS

The apnea–hypopnea index (AHI) for the three subject groups were: moderate/severe OSAS, 21.6 ± 20.5 events/hr (mean ± SD); mild OSAS, 2.1 ± 1.3 events/hr; and normal subjects, 0.2 ± 0.3 events/hr (Table 1). Mean heart rate was not appreciably different between the control group and mild OSAS group and was significantly increased in the moderate/severe OSAS group (Table 1). NL was significantly increased during REM sleep compared to NREM sleep in ~ 88% of children studied regardless of OSAS severity (Figure 2A). Importantly, not only did all subject groups show increases in mean NL from NREM to REM sleep (*p*< 0.001), but the increases were significantly larger in OSAS compared to the normal group (Figure 2A). Linear mixed-effects model analysis of the data for all subject groups and sleep states revealed significant interaction effects between OSAS severity and sleep state for NL (*p* =0.039, Table 2). Thus, although the changes in NL with OSAS levels were quite variable among subjects, NL was a significant predictor for OSAS when interaction with sleep state was taken into account. Interestingly, SampEN also showed significant difference between REM and NREM sleep and significant interaction effects between OSAS severity and sleep state (Table 2). However, linear mixedeffects model analysis showed that the interaction effects were caused by opposing influences of REM sleep and OSAS levels on SampEN without consistent independent effects of OSAS levels *per se* (Table 2).

**Figure 2:**
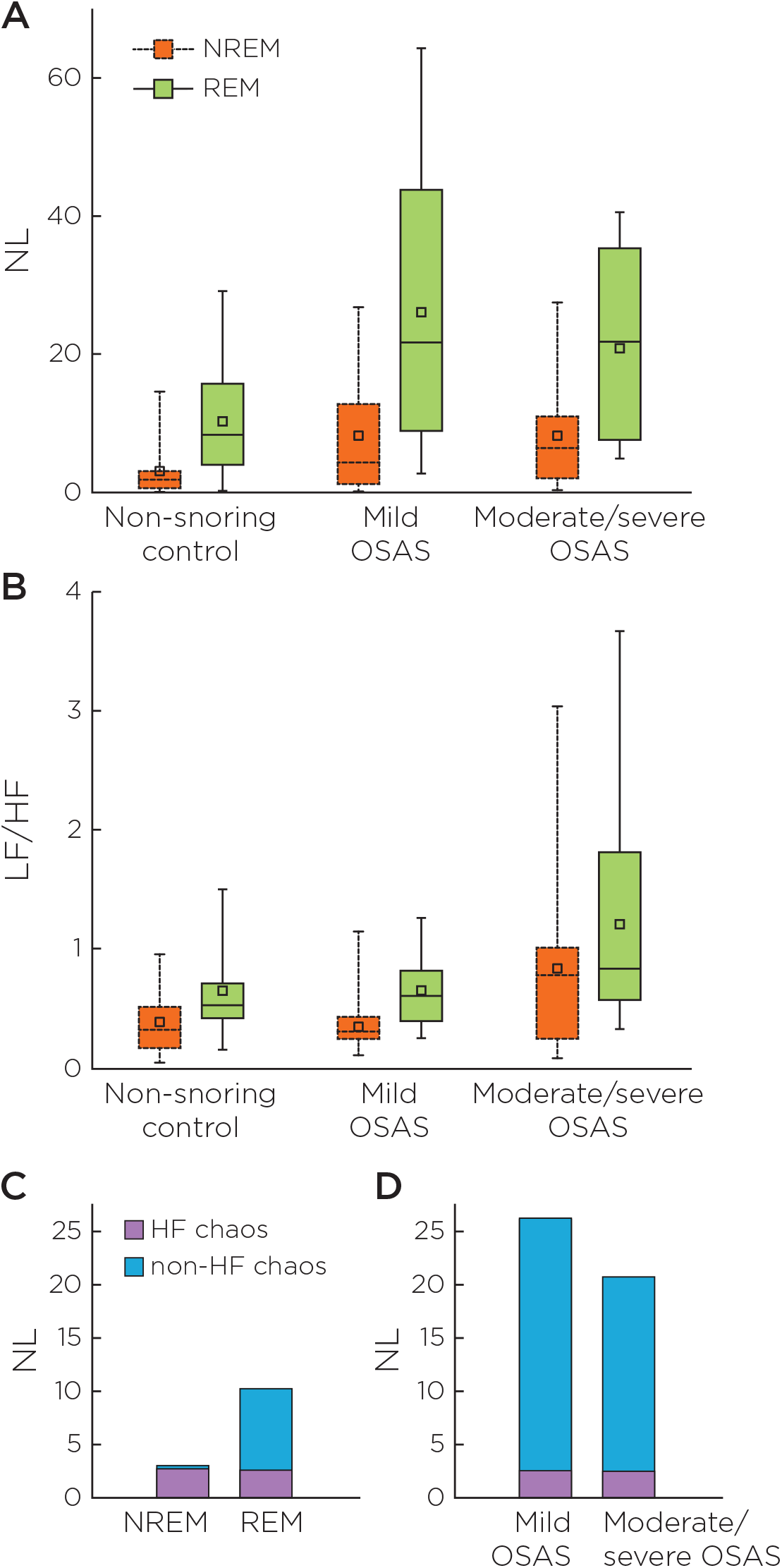
Non-high frequency chaos revealed by combined noise titration and spectral analyses of HRV. (A) Noise limit (NL), a measure of relative chaotic strength (see Supplementary Information); and (B) low- to high-frequency power ratio (LF/HF). The boxes encompass the interquartile ranges with the medians indicated by horizontal divider lines. The whiskers delimit the 5^th^ to 95^th^ percentile of the data distributions. Square markers denote corresponding mean values. Increases in non-HF chaos in REM sleep and OSAS are indicated by concomitant increases in both NL and LF/HF, with corresponding decreases or no change in the HF component (Table 2). Data in Table 2 are summarized by the bar charts in the bottom panels depicting: (C) relative increase in non-HF chaos from NREM to REM sleep in the normal group; and (D) relative increases in non-HF chaos during REM sleep in the mild and moderate/severe OSAS groups compared to the normal group in (c). As a first approximation, non-HF contributions to NL in NREM sleep are assumed small in the normal group and HF chaos contributions to NL are assumed proportional to mean ln(HF) in all conditions.

**Table 2:** Linear mixed-effects model (*N* = 52)

| HRV metric | Sleep state |  | OSAS level |  |  | Interaction |
| --- | --- | --- | --- | --- | --- | --- |
| | REM – NREM (%) | $p$ | Mild OSAS – control (%) | Moderate/severe OSAS – control (%) | $p$ | $p$ |
| SDNN (ms) | –3.5 | 0.244 | –7.6 | –8.8 | 0.719 | NS |
| SDANN (ms) | –7.9 | 0.247 | –11.1 | –0.9 | 0.702 | NS |
| RMSSD (ms) | –11.9 | 0.004 * | –10.4 | –23.2 | 0.641 | NS |
| pNN50 (%) | –32.6 | < 0.001 * | –17.2 | –32.3 | 0.373 | NS |
| Triangular index | –18.0 | < 0.001 * | –20.1 | –3.3 | 0.273 | NS |
| $\ln(\text{LF})$ ($\ln \text{ms}^2$) | 2.6 | 0.084 | 0.0 | 1.9 | 0.750 | NS |
| $\ln(\text{LF})$ ($\ln \text{ms}^2$) | –5.1 | < 0.001 * | –0.0 | –6.7 | 0.510 | NS |
| LF/HF | 62.3 | 0.001 * | –11.2 | 115.7 | < 0.001 * | NS |
| SampEn | –5.4 | 0.017 * | –0.8 | 4.0 | 0.229 | 0.006 * |
| NL | 202.6 | < 0.001 * | 136.7 | 113.5 | 0.314 | 0.039 * |

Data are expressed as % change from control.
\* Statistically significant at the 5% level. NS, not significant.

### B. LF/HF is Increased in REM Sleep and OSAS

None of the HRV metrics except ln(NL) and SampEn showed significant interaction effects between OSAS severity and sleep state (Table 2). Adjusting for OSAS severity, LF/HF was significantly increased in REM sleep compared with NREM sleep (*p* =0.001; also see Figure 2B). The increase in LF/HF reflects primarily a decrease in ln(HF) (*p*< 0.001) indicating parasympathetic withdrawal, with no change in ln(LF). LF/HF was also affected by OSAS although corresponding effects on ln(HF) and ln(LF) were not discernible (Table 2 and Figure 2B); thus LF/HF is a more sensitive indicator of cardiac-autonomic irregularity than the HF or LF component alone. Adjusting for sleep states, LF/HF was significantly higher in moderate/severe OSAS patients compared with normal subjects (*p*< 0.001), but not in mild OSAS patients compared with normal subjects. None of the time-domain metrics (SDNN, SDANN, RMSSD, pNN50, triangular index) were significant predictors for OSAS although some of them were significantly affected by sleep states (Table 2).

### C. Non-HF Chaos Detects Even Mild OSAS

The concomitant increases in NL and LF/HF in REM sleep and OSAS (Figure 2A, B) with corresponding decreases or no change in ln(HF) (Table 2) suggest that the increases in NL in these conditions were not correlated to the parasympathetic-mediated HF component (HF chaos). Accordingly, we hypothesized that such non-HF chaos (Figure 2C, D) may reveal changing cardiac sympathetic-parasympathetic activities that are not discernible by conventional HRV analyses. To determine whether such non-HF chaos may detect OSAS independent of polysomnogram data, we re-evaluated the HRV metrics for all subjects over the full 3-hour segments without separating into different sleep stages (Table 3). Only NL and LF/HF calculated in this manner showed significant differences between the moderate/severe OSAS group and the control group. NL is the only metric that achieved statistical significance (*p*< 0.05) between the mild OSAS group and the control group. ROC curves for the three subject groups showed that NL had higher discriminatory power than LF/HF particularly between the normal and mild OSAS groups (Figure 3 and Table 4).

**Figure 3:**
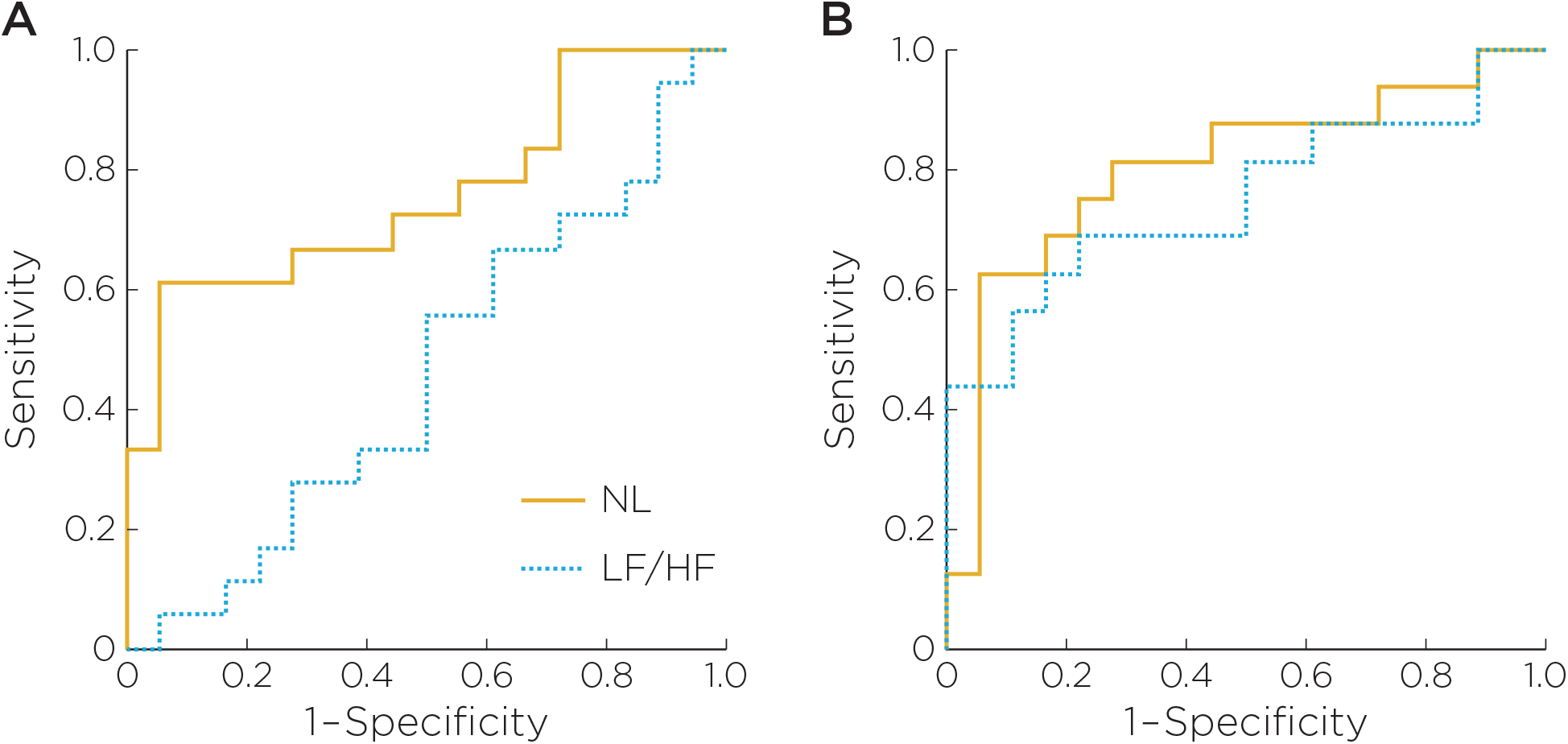
Receiver operating characteristics (ROC) for discrimination between (A) mild OSAS and non-snoring controls; (B) moderate/severe OSAS and non-snoring controls. Curves were derived using noise limit (NL, solid line) and low- to high-frequency power ratio (LF/HF, dotted line) as predictors.

**Table 3:** Group statistics for HRV metrics

| HRV metric | Non-snoring control | Mild OSAS | Moderate/severe OSAS |
| --- | --- | --- | --- |
| SDNN (ms) | 101.5 ± 47.1 | 93.7 ± 28.1 | 92.5 ± 50.3 |
| SDANN (ms) | 44.1 ± 15.3 | 39.2 ± 14.0 | 43.7 ± 20.1 |
| RMSSD (ms) | 101.2 ± 70.0 | 90.7 ± 46.7 | 77.7 ± 59.1 |
| pNN50 (%) | 43.3 ± 25.3 | 35.8 ± 17.3 | 29.3 ± 24.5 |
| Triangular index | 18.3 ± 6.7 | 14.6 ± 3.5 | 17.7 ± 10.2 |
| LF ($\text{ms}^2$) | 1629.0 ± 1410.3 | 1208.9 ± 619.6 | 1600.6 ± 1277.1 |
| HF ($\text{ms}^2$) | 6104.3 ± 7388.4 | 4062.6 ± 3387.4 | 4424.0 ± 6368.6 |
| $\ln(\text{LF})$ ($\ln \text{ms}^2$) | 7.0 ± 1.1 | 7.0 ± 0.5 | 7.1 ± 0.8 |
| $\ln(\text{HF})$ ($\ln \text{ms}^2$) | 8.0 ± 1.4 | 8.0 ± 0.9 | 7.4 ± 1.4 |
| $\text{LF}/\text{HF}$ | 0.50 ± 0.30 | 0.45 ± 0.23 | 1.1 ± 0.67 * |
| SampEn | 0.62 ± 0.07 | 0.61 ± 0.07 | 0.64 ± 0.09 |
| NL | 6.0 ± 4.5 | 14.3 ± 9.3 * | 12.9 ± 7.2 * |

Data (means ± SD) are for full 3-hr recording with both REM and NREM sleep included.
\* Indicates significant difference compared to non-snoring controls ( $p < 0.05$ ).

**Table 4:** OSAS detection performance of NL and LF/HF

| Classification | HRV metric | AUC | Sensitivity (%) | Specificity (%) | Positive predictivity (%) | Negative predictivity (%) |
| --- | --- | --- | --- | --- | --- | --- |
| Mild OSAS vs. control | NL | 0.76 | 61.1 | 94.4 | 91.7 | 70.8 |
|  | LF/HF | 0.46 | 55.6 | 50.0 | 50.0 | 50.0 |
| Moderate/severe OSAS vs. control | NL | 0.80 | 81.3 | 72.2 | 72.2 | 81.3 |
|  | LF/HF | 0.75 | 68.8 | 77.8 | 73.3 | 73.7 |

AUC, area under the ROC curve.

## IV. Discussion

The present study provides the first evidence that non-HF chaos may reveal changes in cardiac sympathetic-parasympathetic activities that are not discernible by conventional HRV metrics alone. Specifically, the increases in NL with concomitant decreases (or no change) in HF power and/or increases in LF/HF suggest that such non-HF chaos is not correlated with the HF component or respiratory sinus arrhythmia. Furthermore, although NL and LF/HF were both significantly elevated over mixed sleep stages in children with moderate/severe OSAS (AHI > 5 /hr), only NL was able to differentiate between mild OSAS (AHI < 5 /hr) and normal controls (AHI < 1 /hr) with demonstrable sensitivity, specificity, positive predictivity, and negative predictivity (Table 4). The univariate ROC performance of non-HF chaos in identifying children with mild OSAS compares favorably to previous classifications of predominantly moderate/severe OSAS patients based on HRV [39, 40] or pulse transit time analyses [29, 41, 42]. The unique ability of the present non-HF heart rate chaos technique to identify children with even mild OSAS paves the way for early screening and intervention of the condition.

Although the present subject populations spanned a wide pediatric age range (1–16 yr) with control subjects being slightly older (by ~ 3.3 yr on average) than OSAS patients, we believe this had little bearing on the study outcome for several reasons. Of note, established normal ranges of HRV during infancy and childhood demonstrate significant decreases in LF/HF during the first year of life but otherwise little changes between 1–15 yr [43], an age range similar to that used in the present study. Although maturation generally has greater effects on other timeand frequency-domain HRV metrics than LF/HF [43, 44], none of these metrics distinguished themoderate/severe OSAS group from the control group as did LF/HF (Tables 2 and 3). The time-and frequency-domain HRV measures presently reported in pediatric subjects with mild and moderate/severe OSAS (average ages of 4.7 ± 3.5 and 4.8 ± 2.2 yr) are similar to established population-based results for older children (average ages of 9.3, 9.8 and 8.9 yr for control, mild OSAS and moderate OSAS groups) [14, 15] that are comparable in age to our control group (8.0 ±4.8 yr). The consensus from those studies and the present study is that time-and frequency-domain HRV metrics are insensitive to mild OSAS in children regardless of age. These observations taken together suggest that age-dependent factors had little influences on changes in HRV metrics both within and between groups in the present study. Indeed, any age-dependent effects would have obscured the primary age-independent effects equally for all HRV metrics. It is therefore highly remarkable that changes in HRV with mild OSAS were discerned by NL and not other HRV metrics in the face of possible age-dependent variations between groups. Importantly, NL discriminated the OSAS groups mainly through their interaction with REM sleep rather than the associated changes in age-dependent factors or OSAS level *per se* (Table 2). As elaborated below, such synergistic effects of OSAS and REM sleep on NL most likely reflect corresponding increases in sympathetic activity.

Recent studies in adult subjects have shown that normal heart rate chaos (with NL > 0) during nighttime may be attributed in large part to HF chaos [27]. Nocturnal increases in HF power and decreases in LF/HF have been reported in pediatric subjects mainly during NREM sleep, particularly slow-wave sleep [13, 14]. Therefore, the heart rate chaos presently found in children during NREM sleep most likely reflects parasympathetic-mediated HF chaos (Figure 2C), as with the previous model for adults [27]. In contrast, the present data show that NL was on average even higher during REM than NREM sleep. This was despite a corresponding decrease in HF power and increase in LF/HF, in agreement with previous reports in children based on conventional methods of HRV power spectrum analysis [13–15]. Thus, it may be reasonably concluded that such non-HF chaos during REM sleep had a predominantly sympathetic instead of parasympathetic mediation. In support of this notion, REM sleep in adults is known to be associated with intermittent, strong and variable sympathetic bursts [7] causing marked cardiorespiratory disturbances [45, 46]. The latter may contribute to the non-HF chaos in children.

Likewise, the present data show that NL was significantly increased over mixed sleep stages while HF power was virtually unchanged (or decreased [14, 15]) in children with OSAS, a pathological condition that is known to provoke persistent elevation of sympathetic activity in adults [8, 9]. The increases in NL with mild and moderate/severe OSAS *per se* were not statistically significant compared with control because of the variability of the NL values within each subject group with considerable overlaps among different groups. However, the OSAS-dependent increases in NL became significant when interaction with sleep state was taken into account suggesting that the sympathoexcitation during REM sleep was probably exacerbated by OSAS. The synergistic effects of OSAS and REM sleep on NL is consistent with the preponderance of apnea episodes during REM sleep in children with OSAS [6]. Again, such OSAS-dependent increase in non-HF chaos is consistent with a predominantly sympathetic instead of parasympathetic mediation. Adding to this notion, a similar trend of paradoxical intermittent increases in NL in the face of decreased HF power has been reported in patients with congestive heart failure [27], another pathological condition that is characterized by sympathoexcitation and parasympathetic impairment [47, 48]. The strikingly consistent pattern of increases in NL concomitant with attenuated or unchanged HF power in REM sleep, OSAS and congestive heart failure, when taken together, strongly suggest that non-HF chaos is a selective marker of sympathoexcitation in these conditions. By contrast, none of the conventional time-or frequency-domain HRV metrics demonstrated interaction effects of OSAS and REM sleep, and none of them was able to detect mild OSAS in children.

These observations underscore the increasing evidence that HRV is intrinsically a nonlinear phenomenon and cannot be fully deciphered by using conventional time-or frequencydomain linear analyses [49, 50]. On the other hand, current approaches to studying the nonlinear behavior of HRV using classical nonlinear dynamics methods (such as Lyapunov exponent, correlation dimension, surrogate data method, etc.) or statistical physics methods (such as 1/*f* scaling, entropy, mono-or multifractal analysis, etc.) suffer from the lack of sensitivity, specificity, and robustness in discriminating the nonlinearity or complexity of HRV from measurement noise and dynamic noise/intrinsic process disturbance (for a critique of these methods see [51] and Appendix S1 in [27]). The present study showed that a popular statistical physics method (SampEn) was insensitive to OSAS in children, consistent with the reported lack of accuracy of this method in detecting adult OSAS [52]. To circumvent these difficulties, the noise titration technique originally proposed to overcome the effect of measurement noise has been extended to the detection of noise-induced chaos in the presence of dynamic noise (see section “Nonautonomous Nonlinear Dynamical Systems” in [24] and Appendix S1 in [27] and Figure 2 in [51]). This generalized framework [53] has led to the previous identification of HF chaos [27]) and the present identification of non-HF chaos as potential noninvasive markers of state-dependent changes in cardiac parasympathetic–sympathetic activities during REM sleep, non-REM sleep and wakefulness in health and in OSAS.

A potential limitation of the present study is that sympathoexcitation was inferred indirectly from increases in NL assuming unchanged or decreased parasympathetic activity as assessed by ln(HF) and LF/HF. Such non-HF chaos provides only a relative measure of the predicted sympathoexcitation, as the resultant increase in NL may be offset by any concurrent decrease in HF chaos in a nonlinear fashion. This may explain why NL was lower in the moderate/severe OSAS group than the mild OSAS group, as the correspondingly higher LF/HF and mean heart rate (and lower HF power [14, 15]) indicate concurrent parasympathetic withdrawal. Further, the HF component and LF/HF are known to be influenced also by changes in breathing pattern, which may potentially confound their interpretation as indices of parasympathetic activity during obstructive apnea episodes [54, 55]. Although the effect of apnea on respiratory–cardiac coupling can in theory be compensated to some extent through nonlinear multivariate modeling involving simultaneous ventilatory and blood pressure measurements [56] (ideally also end-tidal pCO_2_, blood oximetry and esophageal pleural pressure or diaphragmatic EMG recordings), such elaborate cardiorespiratory recordings are beyond the scope of clinical polysomnography and are impracticable for the purpose of home screening test. Despite this, it is important to note that respiratory activity and respiratory sinus arrhythmia typically persist during obstructive apnea episodes even when respiratory airflow is nil, unlike the abolition of both during central apnea [57]. This may explain why ln(HF) remained unchanged or only minimally decreased in the moderate/severe OSAS group despite the presence of pronounced obstructive apnea hypopnea events (see also [14, 15]). In contrast, the effect of apnea episodes on NL is likely to involve more than a direct respiratory–cardiac coupling. Our previous studies in awake subjects have shown that NL is increased during breath-holding even though respiratory sinus arrhythmia is abolished by the ensuing apnea [58, 59]. Such an increase in NL absent HF chaos again likely reflects apnea-induced sympathoexcitation, as demonstrated recently by corresponding increases in muscle sympathetic nerve activity secondary to baroreflex and chemoreflex activations during breath-holding maneuvers [60]. This suggests that non-HF chaos is a reliable index of sympathoexcitation in these conditions despite possible disruption of the HF component during apnea episodes. Further studies are needed to elucidate the independent effects of breathing pattern and other factors on HF chaos and non-HF chaos during REM sleep, non-REM sleep and wakefulness in health and in OSAS.

## V. Conclusion

We have provided strong evidence in support of the hypothesis that non-HF chaos is a selective noninvasive marker of the sympathoexcitation associated with REM sleep, OSAS and other conditions. The presently demonstrated sensitivity and specificity of non-HF chaos in tracking REM sleep- and OSAS-dependent autonomic abnormalities open the possibility of a home screening test of sympathetic-parasympathetic imbalance for early diagnosis of pediatric and adult OSAS [5, 20] and a spectrum of cardiovascular diseases [61]. Because autonomic function is influenced by many factors and could vary considerably with time and between subjects, rigorous experimental designs and data analyses are critical for optimal test performance. Future studies will further evaluate and refine this new technique as a noninvasive probe of changing sympathetic–parasympathetic activities in a larger cohort of pediatric and adult patients with OSAS and other autonomic disorders in the home and laboratory environments.

## Acknowledgments

This work was supported by National Institutes of Health grants K23-HL073238 (EK), R01HL058585 (CLM), R21-HL075014, R01-HL079503, R01-HL067966 and RC1-RR028241 (CSP).

## Supplemental Information

### Noise Titration Technique

The heart of the noise-titration algorithm [24] is the Volterra autoregressive series (VAR) nonlinearity detection algorithm. Given an *N*-point time series *x*(*n*)=[*x*(1),*x*(2),…,*x*(*N*)], the Volterra series expansion formulates the input/output relationship as a polynomial combination of input delays. Within this framework, we analyze the univariate time series by using a discrete VAR series of degree *d*, and memory *κ*, as a model to calculate the predicted time series [25]:

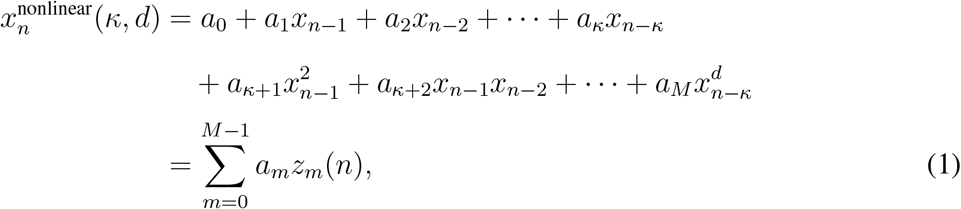

where the functional basis {*z_m_*(*n*)} is composed of all the distinct combinations of the embedding space coordinates (*x*_*n*−1_,*x*_*n*−2_,…,*x*_*n*−*κ*_) up to degree *d*, with a total dimension 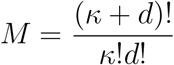; and {*a_m_*(*n*)} is the set of Volterra kernel functions, characterizing the autoregressive behavior of the dynamical system. Each model is parameterized by *κ*, the embedding dimension, and *d*, the degree of the nonlinearity of the model (i.e. *d* =1 for linear model and *d*> 1 for nonlinear model). The coefficients *a_m_* are recursively estimated by using the Korenberg fast orthogonal algorithm [65].

The goodness of fit of a model (linear vs. nonlinear) is measured by the normalized residual sum-of-square errors

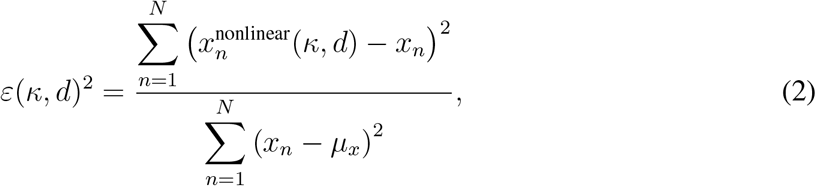

where 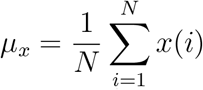 and *∊*(*κ*,*d*)^2^ is in effect a normalized variance of the residual error. The optimal model {*κ*_optimal_,*d*_optimal_} is the one that minimizes the Akaike cost function

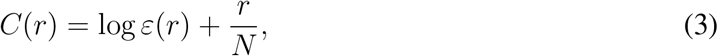

where *r* ∈ [1,*M*] is the number of polynomial terms of the truncated Volterra expansion. The Akaike information criterion measures the balance between the goodness of fit of a model and the model complexity (parsimony principle). In addition to linear vs. nonlinear hypothesis testing, the VAR method provides a sufficient test for chaotic dynamics when used in conjunction with a numerical “noise titration” procedure: the dynamics of nonlinearity are tested on the time series. If linearity is detected, then the noise titration method rejects the null hypothesis of no chaotic behavior. If nonlinearity is detected, small amounts of Gaussian noise (1% of signal power) are successively added until nonlinearity is no longer detected (within a prescribed level of statistical confidence). The maximum noise added before nonlinearity goes undetected is known as the noise limit (NL). Under the noise titration scheme, a noise limit greater than zero represents the detection of chaotic dynamics. In addition, the noise limit mirrors the maximal Lyapunov exponent of the system dynamics [24].

